# Preclinical validation of AAV9-TECPR2 gene therapy in a novel knock-in model of TECPR2-related disorder

**DOI:** 10.64898/2026.03.02.708636

**Authors:** Bruna Lenfers Turnes, Peter Casey-Caplan, Leo Mejia, Tiffany Berry, Julie Zhao, Francesco Villa, Emma Cropper, Maryam Arab, Biyao Zhang, Darred Surin, Darius Ebrahimi-Fakhari, Silmara de Lima, Alan S. Kopin, Nick Andrews, Nathaniel Hodgson, Michela Fagiolini

## Abstract

*TECPR2*-related disorder is a rare, autosomal recessive neurodevelopmental and neurodegenerative disease characterized by early-onset motor dysfunction, sensory- and autonomic neuropathy, and progressive neurological decline with early mortality. Currently, there are no effective treatments for individuals affected by this debilitating condition. To advance our understanding of disease mechanisms and explore therapeutic strategies, we developed and then characterized a knock-in (KI) mouse model carrying the human *TECPR2* c.1319delC frameshift mutation.

TECPR2-KI mice exhibit a subset of disease-relevant phenotypes, most prominently abnormal gait, along with reduced body weight and altered tactile sensitivity. We additionally observe a reduction in acoustic startle responses, consistent with dysfunction of brainstem-associated sensorimotor pathways. Histopathological analyses reveal progressive accumulation of axonal spheroids in the dorsal column nuclei, together with abnormalities in autophagy-related markers, features previously reported in individuals with *TECPR2*-related disorder.

To assess the therapeutic potential of gene replacement, we delivered *TECPR2* via intracisternal infusion of AAV9/*TECPR2* in neonatal KI mice. Gene therapy restored mechanosensory function, normalized gait and startle responses, maintain autophagic homeostasis, and partially reduced axonal pathology. These findings demonstrate that *TECPR2*-associated deficits are not only replicable in this new mouse model but are also amenable to postnatal intervention.

Our study introduces a genetically accurate murine model of TECPR2 deficiency, identifies brainstem-associated phenotypes, and provides preliminary evidence supporting the feasibility of AAV9-mediated TECPR2 gene delivery, establishing a foundation for future translational research in a currently untreatable disease.

**One-Sentence Key Message:** TECPR2 deficiency disrupts brainstem sensory–motor circuits, impairing autophagy and tactile, gait, and startle function and is prevented by neonatal AAV9.

## INTRODUCTION

*TECPR2*-related disorder is a devastating, ultra-rare neurodevelopmental and neurodegenerative condition for which no disease-modifying therapies currently exist. It is classified along the spectrum of the hereditary sensory and autonomic neuropathies (specifically HSAN9, OMIM #615031) and the complex hereditary spastic paraplegias (HSPs). Since its first identification in 2012, fewer than 40 individuals have been reported worldwide, underscoring both the rarity and the limited clinical awareness of the disorder (1-5).

*TECPR2*-related disorder is caused by biallelic loss-of-function mutations in the *TECPR2* gene, which encodes the tectonin beta-propeller repeat-containing protein 2, a multifunctional scaffold protein with broad expression across the central nervous system (CNS). TECPR2 has been implicated as a key regulator of the autophagy–lysosome pathway, particularly in mediating the targeting of autophagosomes to lysosomes and promoting autophagic flux (6). Efficient autophagic clearance is essential for maintaining neuronal health, especially in long-projecting neurons that depend on the constant turnover of organelles and proteins to preserve axonal integrity (7,8). Accordingly, dysfunction of autophagy-related genes has been increasingly associated with a wide range of neurodevelopmental and neurodegenerative disorders (9-12). Loss of TECPR2 function results in impaired autolysosomal fusion, accumulation of protein aggregates, and formation of neuroaxonal spheroids, ultimately leading to axonal dystrophy and neuronal loss (6).

Pathological spheroids have been documented not only in affected human patients but also in other species, highlighting the conserved nature of TECPR2-deficiency related axonopathy. For instance, Spanish Water Dogs with a spontaneous *TECPR2* loss-of-function mutations display progressive axonal degeneration, and a CRISPR-Cas9-generated *Tecpr2* knockout mouse model exhibits comparable neuropathological features (11,5). These findings reinforce the importance of developing robust and genetically faithful animal models to study disease mechanisms and evaluate potential interventions.

Clinically, *TECPR2*-related disorder manifests with global developmental delay and later intellectual disability, early-onset hypotonia, sensory and autonomic neuropathy, progressive spasticity, and progressive bulbar dysfunction with dysphagia and central apneas. Dysfunction of brainstem-mediated circuits, contributing to dysphagia, impaired reflexes, and central apneas, accounts for significant morbidity and early mortality. Most affected individuals succumb to disease in the first or second decade of life, and survivors experience profound, lifelong disability (1,3,5). Despite the recognition of these hallmark features, the precise temporal progression of neuropathological changes and their mechanistic underpinnings remain poorly defined, representing a major barrier to therapeutic development.

Efforts to identify therapeutic strategies for *TECPR2*-related disorders have been hindered by the rarity of the condition and the lack of suitable *in vivo* models that capture disease-relevant mutations and phenotypes. Although a recent study demonstrated that antisense oligonucleotides (ASOs) targeting exon 8 of TECPR2 produced promising effects *in vitro* (13), the development of candidate therapeutics for *TECPR2*-related disorders, genetic approaches (e.g. ASO, gene therapy) or otherwise, will require *in vivo* models to test the efficacy of the approach.

Given these limitations, gene replacement therapy using adeno-associated virus serotype 9 (AAV9) has emerged as a potential alternative. AAV9 vectors are well characterized for their ability to transduce both neurons and glia across the CNS following intracisternal, intrathecal or intravenous delivery, and they have been successfully used in the treatment of other pediatric neurological disorders, including spinal muscular atrophy (14). Importantly, this approach allows for full-length *TECPR2* gene delivery irrespective of the specific mutation, offering a mutation-agnostic strategy for therapeutic intervention.

To address the critical gap in preclinical models, we generated a knock-in (KI) mouse line carrying the human *TECPR2* c.1319delC frameshift mutation. This model enables the assessment of behavioral, sensory, and neuropathological alterations associated with TECPR2 deficiency and was used to evaluate intracisternal AAV9-mediated *TECPR2* gene delivery at neonatal stages. Together, this work provides (i) a genetically accurate *in vivo* model for investigating *TECPR2*-related pathophysiology, (ii) insight into previously unrecognized sensorimotor deficits involving brainstem circuits, and (iii) initial preclinical evidence supporting the feasibility of *TECPR2* gene replacement.

## RESULTS

### Behavioral alterations in *TECPR2*-KI mice recapitulate aspects of the clinical phenotype observed in *TECPR2*-related disorder

To model human *TECPR2*-associated disease, we generated a homozygous *Tecpr2* c.1319delC knock-in (KI) mouse line. This mutation introduces a premature stop codon at the same position as the pathogenic c.1319delT founder variant identified in a large cohort of patients (5) thereby recapitulating the genetic basis of disease. We performed longitudinal behavioral phenotyping at postnatal days 30, 60, and 90 to assess developmental and progressive manifestations.

At P90, *TECPR2*-KI mice exhibited a small significant reduction in body weight compared with WT littermates (Fig. 1A), consistent with reports of growth impairment in some individuals with *TECPR2*-related disorder (5). We examined their self-grooming behavior over a 10-minute video recording with automatic classification of behaviors and observed that TECPR2-KI mice show increased grooming behavior at P90 (Fig. 1B). Given reports of sensory abnormalities in patients, we assessed tactile sensitivity in *TECPR2*-KI mice using von Frey filaments. At >P90, *TECPR2*-KI mice exhibited a small increase in mechanical thresholds compared with WT controls (Fig. 1C), while responses to pin prick and noxious heat were preserved (data not shown). Together, these findings suggest subtle and age-dependent alterations in tactile sensitivity in *TECPR2*-KI mice, though the magnitude and consistency of the effect warrant cautious interpretation.

**Figure 1:**
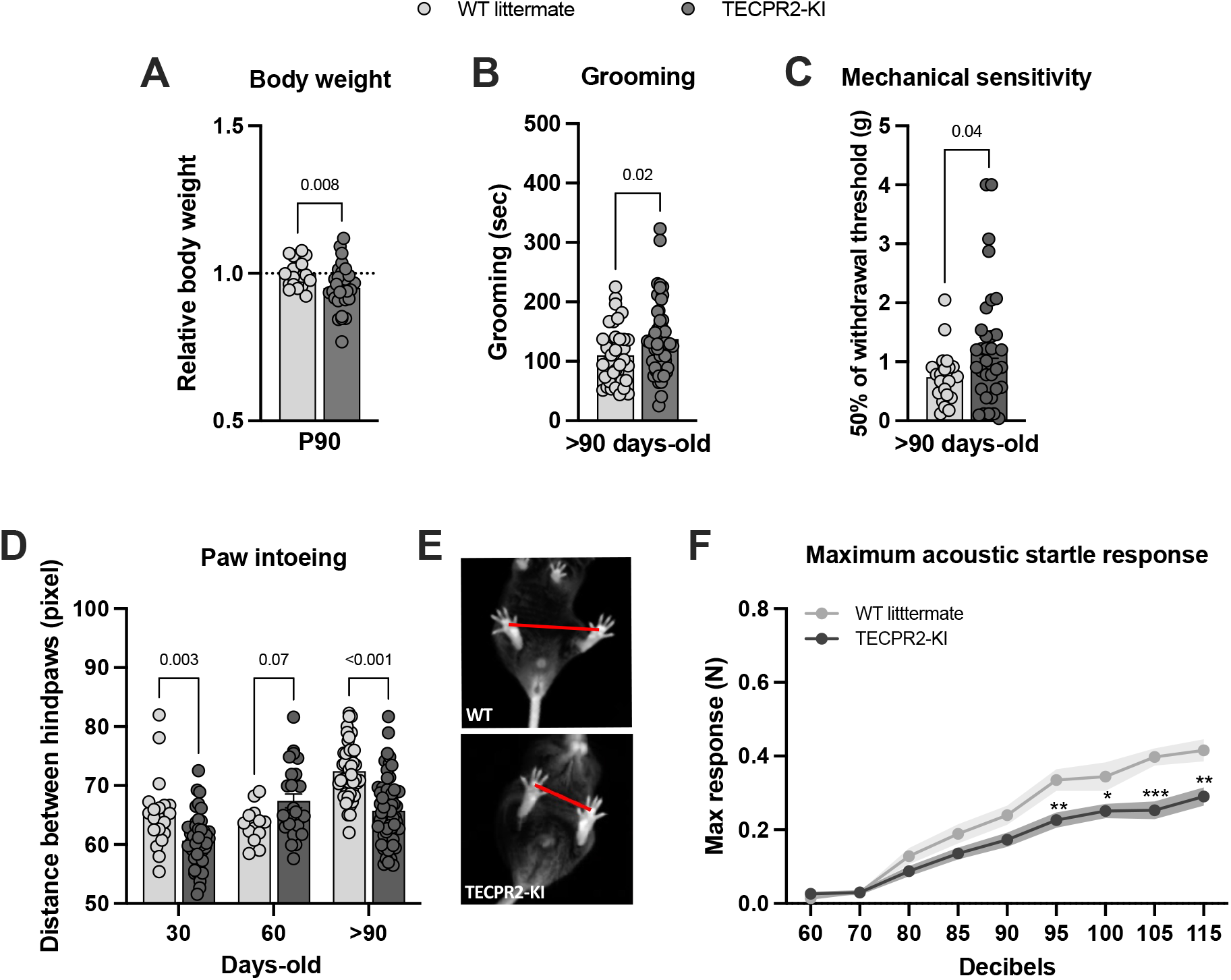
*TECPR2*-KI mutation [NM_001081057.2; c.1319delC, p.Ser440Serfs19*] leads to changes in behavior phenotype. **A**, Relative body weight of *TECPR2*-KI and WT littermate male and female mice at P90 (*n* = 20-29 mice per group, data are presented as mean values ± s.e.m., *P* values from unpaired t test). **B**, Grooming time of *TECPR2*-KI and WT littermate male and female mice at >P90 (*n* = 23-41 mice per group; data are presented as mean values ± s.e.m., *P* values from unpaired t test). **C**, Mechanical withdrawal thresholds of *TECPR2*-KI and WT littermate male and female mice using an up–down von Frey method at >P90 (*n* = 21-33 mice per group; data are presented as mean values ± s.e.m., *P* values from unpaired t test). **D**, Distance between hindpaws used as a measure of intoeing in male and female *TECPR2*-KI and WT littermates over time (*n* = 14-62 mice per group, data are presented as mean values ± s.e.m., *P* values from two-way ANOVA followed by Šídák’s multiple comparisons test). **E**, Representative images of intoeing in *TECPR2*-KI and WT littermate control. **F**, Maximum acoustic startle response in male and female *TECPR2*-KI and WT littermates at P90 (*n* = 16-18 mice per group, data are presented as mean values ± s.e.m., *\*P < 0*.*03, **P < 0*.*002, ***P < 0*.*001* from two-way ANOVA followed by Šídák’s multiple comparisons test).

Given the motor impairment present in patients with *TECPR2*-related disorder, we also analyzed gait dynamics using a novel bottom-up imaging and data acquisition technology and analysis platform (19). *TECPR2*-KI mice displayed progressive gait abnormalities, characterized by a reduced hindlimb base width (increased intoeing), starting as early as P30 and worsening with age (>P90) (Fig. 1D,E), supporting a relatively early behavioral onset of defects. Brainstem nuclei implicated in tactile processing also contribute to sensorimotor integration of auditory inputs. Therefore, we evaluated the acoustic startle reflex and observed that *TECPR2*-KI mice exhibited a markedly reduced maximum startle amplitude at >P90 compared with WT littermates (Fig. 1F). This represents a previously unrecognized phenotype in *TECPR2*-related disease and suggests an additional brainstem-mediated sensory–motor impairment. Notably, grip strength and locomotion did not differ between genotypes (Supplementary Fig. 1A,B), excluding generalized muscle weakness as a cause of gait or startle abnormalities.

Collectively, these findings indicate that *TECPR2*-KI mice exhibit a subset of disease-relevant phenotypes, most prominently brainstem acoustic startle disfunction, progressive gait abnormalities, along with additional mechanosensory deficits.

### *TECPR2*-KI mice develop progressive axonal pathology and defective autophagy in the brainstem

Histopathological examination revealed that *TECPR2*-KI mice, but not WT littermates, developed progressive axonal spheroid accumulation in the gracile and cuneate nuclei of the medulla oblongata (Fig. 2A). These spheroids, representing focal axonal swellings, were also present in the cervical–thoracic spinal cord (data not shown) but not in cortical layers, indicating selective vulnerability of brainstem and spinal sensory tracts. Quantification demonstrated that spheroid accumulation increased with age, first appearing at P60 (∼100 spheroids per mm^2^) and becoming more pronounced at P90 (∼300 spheroids per mm^2^) (Fig. 2B). This progressive axonal pathology closely mirrors previous reports in *Tecpr2* knockout mice (6) and parallels the neurodegeneration observed in patients. Ultrastructural analysis by transmission electron microscopy revealed prominent structural abnormalities in *TECPR2*-KI mouse brainstem, showing atypical autophagosome morphology and robust accumulations of single- and double-membrane-limited autophagosome vacuoles with electron-dense content, consistent with impaired autophagosome clearance (Supplementary Fig. 3A–D). Supporting this, RT–qPCR confirmed reduced endogenous Tecpr2 expression in KI brainstem tissue (Fig. 2C). Immunostaining showed significant accumulation of the autophagy substrate SQSTM1/p62 in mutant mice, while levels of neurofilament heavy chain (NF-H), a biomarker of axonal integrity, were unchanged (Fig. 2D–F). These findings indicate that loss of TECPR2 function causes selective autophagic defects in brainstem nuclei, leading to progressive spheroid formation and axonal pathology.

**Figure 2.**
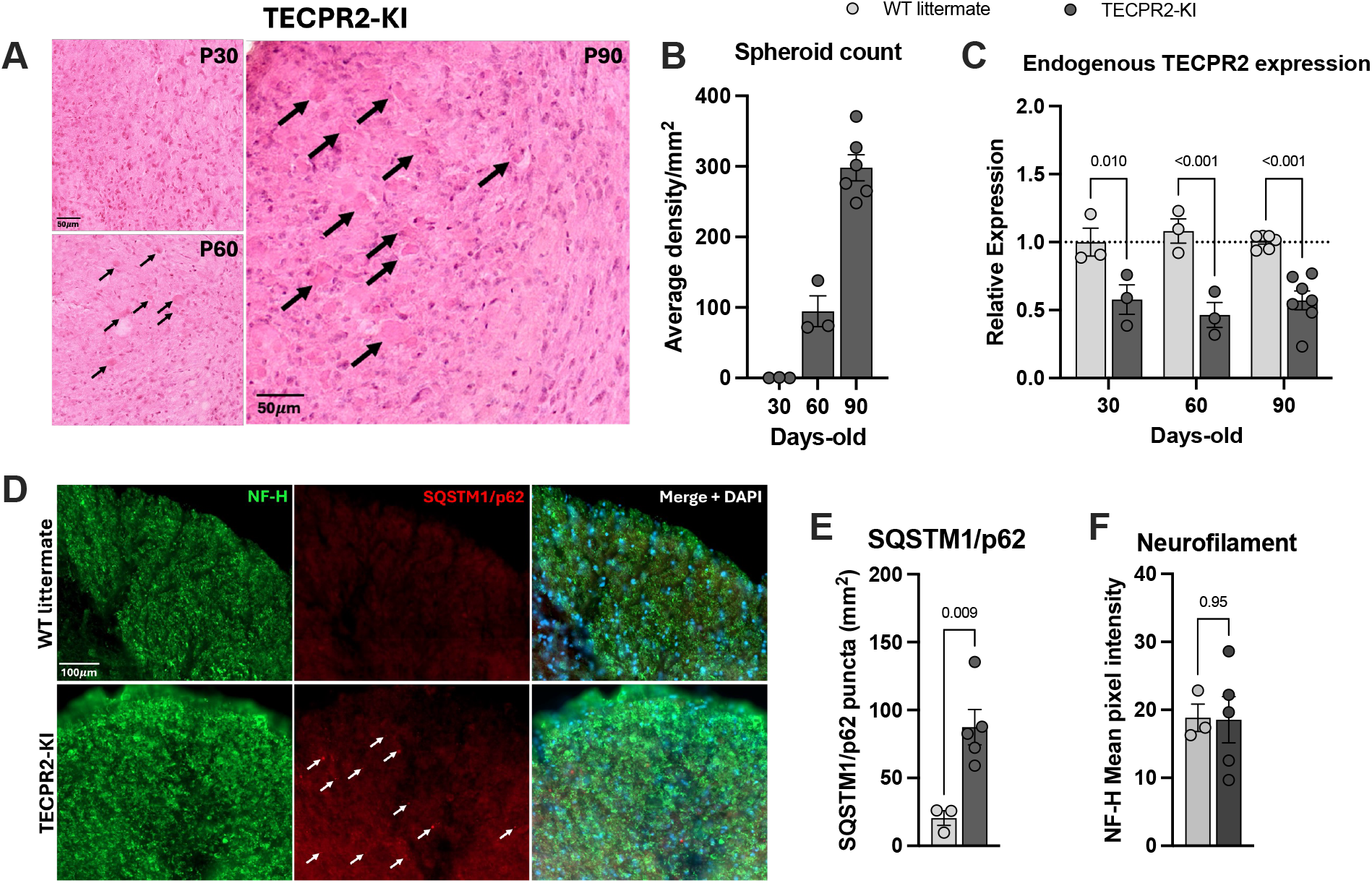
*TECPR2*-KI mutation induces accumulation of spheroids and SQSMT1/p62 in the brainstem. **A**, Representative H&E staining of coronal brainstem sections (cuneate and gracile nuclei) of *TECPR2*-KI mice showing (indicated by arrows) progressive appearance of spheroids at P30, P60 and P90. Scale bars, 50 μm. **B**, Quantification of spheroids in brainstem of *TECPR2*-KI mice (*n* = 3-6 mice per group). **C**, RT–qPCR quantification of endogenous TECPR2 in brainstem of *TECPR2*-KI and WT littermates male and female mice (*n* = 3-7 mice per group, data are presented as mean values ± s.e.m., *P* values from two-way ANOVA followed by Šídák’s multiple comparisons test). **D**, Representative SQSTM1/p62 and neurofilament heavy chain (NF-H) staining of brainstem of TECPR2-KI and WT littermates at P90, showing accumulation of SQSMT1/p62, general marker of autophagic flux, in TECPR2-KI mice. Scale bars, 100 μm. **E**, Quantification of SQSTM1/p62 and **F**, NF-H (*n* = 3-5 mice per group; data are presented as mean values ± s.e.m., *P* values from unpaired t test).

### AAV9-mediated TECPR2 gene replacement is associated with improvements in select motor and sensory measures

To determine whether gene replacement could mitigate disease phenotypes, we delivered a single intracisternal injection of AAV9.pU1a-TECPR2_WT (VCAV-04485, 5.00E+12 GC/mL; 1uL) to *TECPR2*-KI mice and WT littermates at P1–3 (Fig. 3A). We chose the cisterna magna as the route of administration because of its proximity to the dorsal column nuclei, the earliest site of pathology, and its established efficacy in neonatal CNS gene therapy (14). The treatment was well tolerated in WT mice, supporting safety and long-term tolerability. In KI mice, AAV9-based gene placement prevented only a subset of the disorder manifestations. Although transient prevention of body weight loss was noted in males at P30 (Supplementary Fig. 2A), treated KI mice continued to exhibit reduced body weight at >P90 (Fig. 3B) and excessive grooming behavior (Fig. 3C). Improvements were observed across several functional measures. AAV9/TECPR2-treated KI mice showed normalized mechanical sensitivity at >P90 (Fig. 3D), improved hindlimb base width (paw intoeing) at P30 (Fig. 3E), and complete recovery of acoustic startle response amplitudes to WT levels (Fig. 3F). Together, these findings indicate that AAV9-mediated TECPR2 expression can partially ameliorate mechanosensory and brainstem-associated sensorimotor deficits, while growth and grooming abnormalities remain largely unaffected.

**Figure 3.**
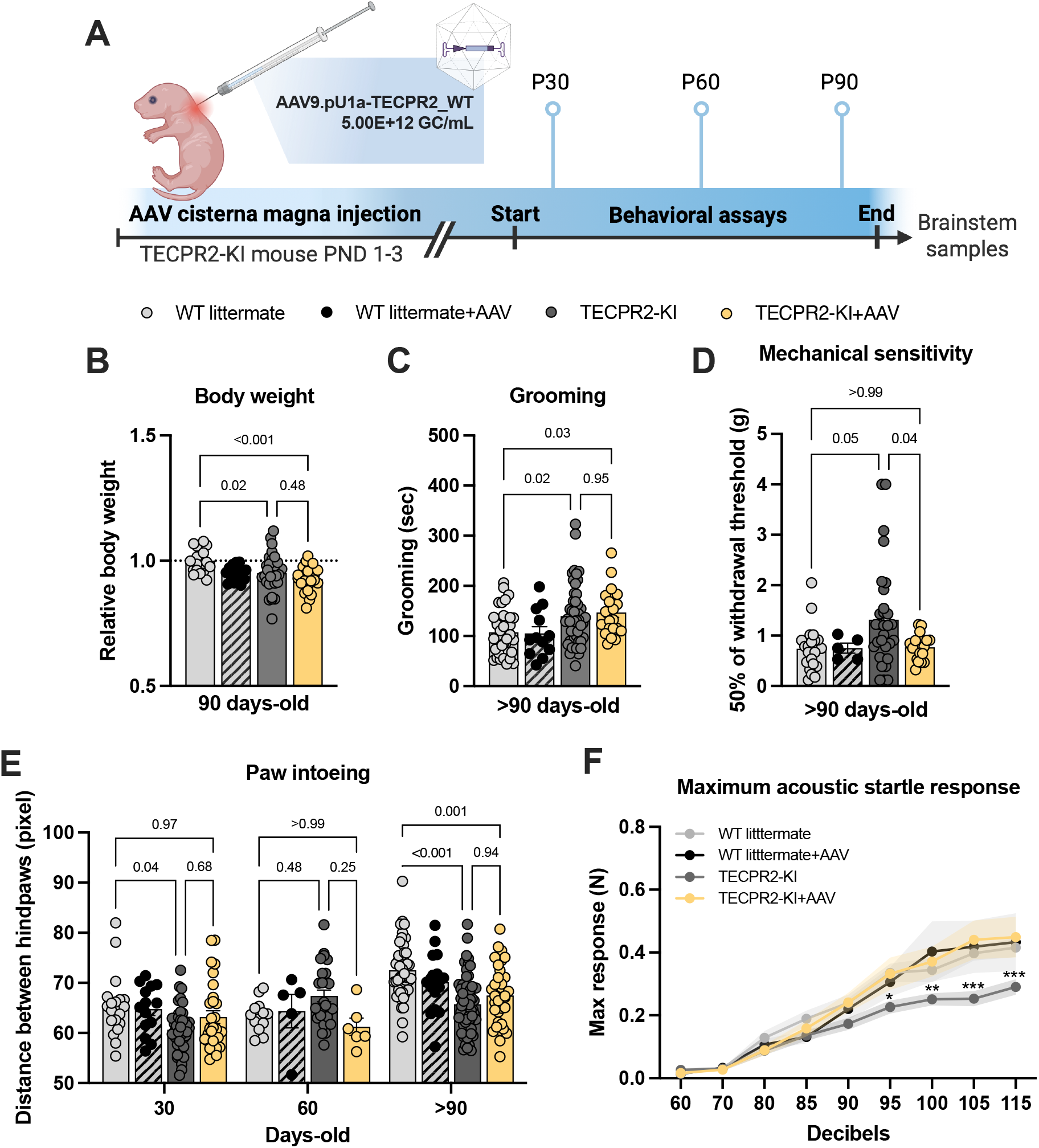
AAV9/TECPR2 mediated gene replacement in *TECPR2*-KI mice is associated with improvements in select motor and sensory measures. **A**, Illustration of experimental regimen. **B**, Relative body weight of TECPR2-KI, WT littermate, TECPR2-KI+AAV and WT littermate + AAV male and female mice at P90 (*n* = 17-29 mice per group, data are presented as mean values ± s.e.m., *P* values from ANOVA followed by Tukey’s multiple comparisons test). **C**, Grooming time of TECPR2-KI, WT littermate, TECPR2-KI+AAV and WT littermate + AAV male and female mice at >P90 (*n* = 12-50 mice per group; data are presented as mean values ± s.e.m., *P* values from ANOVA followed by Tukey’s multiple comparisons test). **D**, Mechanical withdrawal thresholds of *TECPR2*-KI, WT littermate, *TECPR2*-KI+AAV and WT littermate + AAV male and female mice using an up–down von Frey method (*n* = 5-33 mice per group; data are presented as mean values ± s.e.m., *P* values from one-way ANOVA followed by Dunnett’s T3 multiple comparisons test). **E**, Distance between hindpaws used as a measure of intoeing in male and female *TECPR2*-KI, WT littermate, *TECPR2*-KI+AAV and WT littermate + AAV over time (*n* = 5-62 mice per group, data are presented as mean values ± s.e.m., *P* values from two-way ANOVA followed by Šídák’s multiple comparisons test). **F**, Maximum acoustic startle response in male and female *TECPR2*-KI, WT littermate, *TECPR2*-KI+AAV and WT littermate + AAV (*n* = 7-30 mice per group, data are presented as mean values ± s.e.m., *\*P < 0*.*05, **P < 0*.*01, ***P < 0*.*001* from two-way ANOVA followed by Tukey’s multiple comparisons test).

### AAV9-mediated TECPR2 gene placement reduces spheroid pathology and improves autophagic flux

We next examined whether AAV9-TECPR2 treatment could mitigate axonal and autophagic defects in *TECPR2*-KI mice. Histological analysis revealed that intracisternal AAV9 delivery at P1–3 partially prevented spheroid accumulation in the gracile and cuneate nuclei, reducing their density by approximately 30% at P90 compared with untreated mutants (Fig. 4A,B). Robust viral-mediated expression of TECPR2 was confirmed in the brainstem of treated animals (Fig. 4C). Importantly, treated WT littermates displayed normal histoarchitecture and physiological TECPR2 expression, indicating that the therapy was safe and well tolerated. Immunohistochemical analyses further showed improvement of autophagy-related markers in AAV9-treated KI mice. Accumulation of the autophagy substrate SQSTM1/p62, prominent in untreated mutants, was similar to WT in the brainstem of treated animals (Fig. 4D,E), while levels of neurofilament heavy chain (NF-H) were comparable across groups (Fig. 4F).

**Figure 4.**
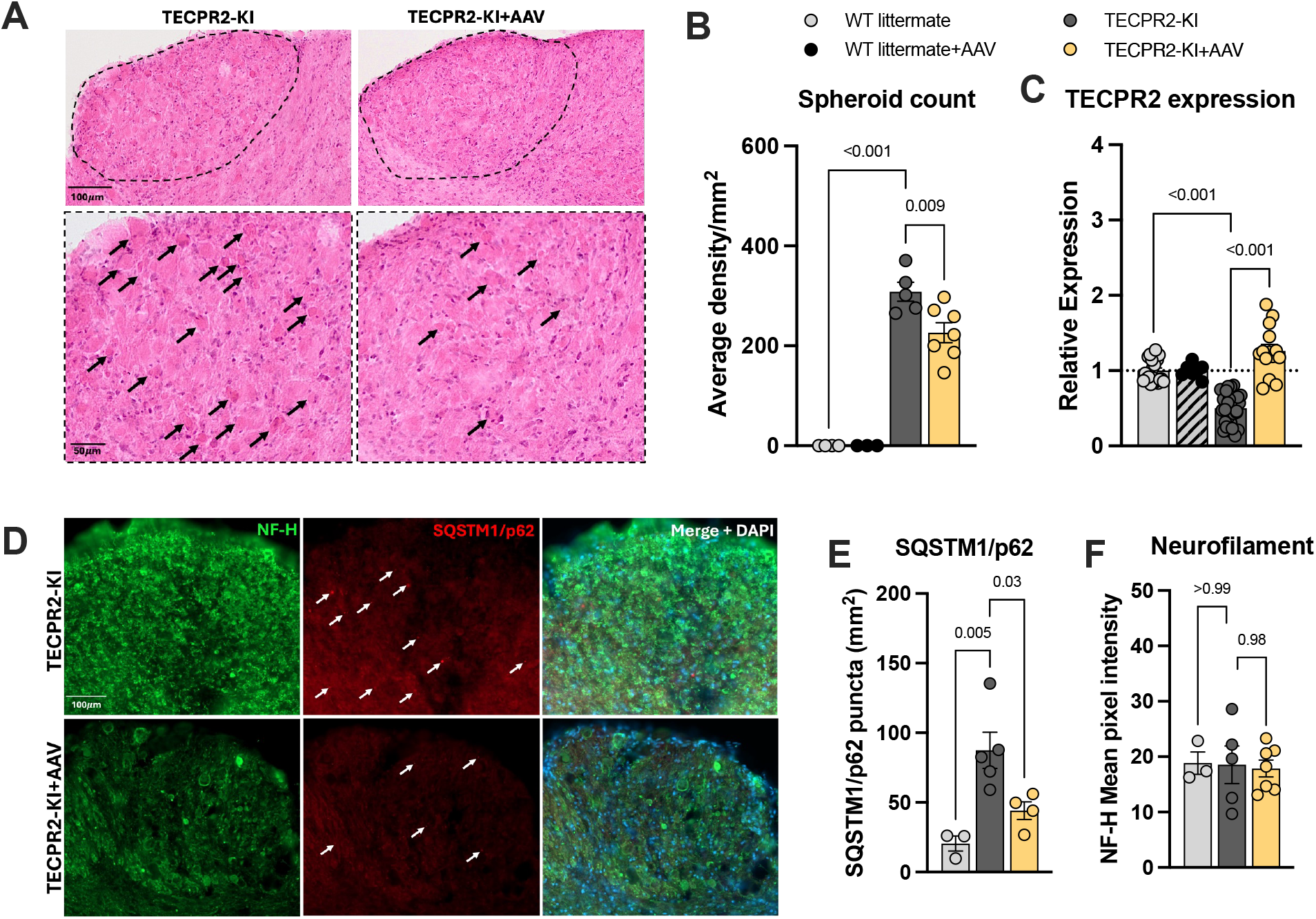
AAV9/TECPR2 mediated gene replacement in *TECPR2*-KI mice reduces spheroid accumulation and ameliorates autophagic flux at P90. **A**, Representative H&E staining of coronal brainstem sections of *TECPR2*-KI, WT littermates and *TECPR2*-KI+AAV mice showing (indicated by arrows) spheroid accumulation at P90. Scale bars, 50 and 100 μm. **B**, Quantification of spheroids in brainstem of *TECPR2*-KI, WT littermate, *TECPR2*-KI+AAV and WT littermate + AAV male and female mice (*n* = 4-7 mice per group, data are presented as mean values ± s.e.m., *P* values from one-way ANOVA followed by Dunnett’s multiple comparisons test). **C**, RT–qPCR quantification of endogenous and viral TECPR2 in brainstem of *TECPR2*-KI and WT littermates (endogenous), and *TECPR2*-KI+AAV and WT littermate + AAV (viral) male and female mice (*n* = 7-30 mice per group, data are presented as mean values ± s.e.m., *P* values from one-way ANOVA followed by Dunnett’s multiple comparisons test). **D**, Representative SQSTM1/p62 staining of brainstem of *TECPR2*-KI, WT littermates and TECPR2-KI+AAV at P90. Scale bars, 100 μm. **E**, Quantification of SQSTM1/p62 and **F**, NF-H in brainstem of TECPR2-KI, WT littermates and TECPR2-KI+AAV male and female mice (*n* = 3-7 mice per group, data are presented as mean values ± s.e.m., *P* values from one-way ANOVA followed by Turkey’s multiple comparison test).

## DISCUSSION

*TECPR2*-related disorder is a rare neurodevelopmental and neurodegenerative disease for which no disease-modifying therapies currently exist. In this study, we describe a knock-in mouse model carrying the recurrent *TECPR2* c.1319delC founder mutation and characterize behavioral, sensory, and neuropathological abnormalities associated with TECPR2 deficiency. Our data suggest an association between impaired autophagic flux and selective involvement of brainstem sensory– motor circuits and identify an acoustic startle deficit as a functional readout relevant to circuit dysfunction. We further evaluate early postnatal AAV9-mediated *TECPR2* gene delivery and observe partial improvements in autophagy-related markers, axonal pathology, and select functional outcomes. Together, these results provide a framework for investigating disease mechanisms and support further evaluation of gene replacement approaches in *TECPR2*-related disorder, particularly to treat brain-stem related disease manifestations which account for early mortality.

Neurodevelopmental and neurodegenerative disorders caused by impaired autophagy present a major translational challenge (11,12) as they often combine early circuit dysfunction with progressive neurodegeneration but lack validated disease models and actionable outcome measures. *TECPR2*-related disorder exemplifies this gap: despite clear genetic causation and severe clinical manifestations, therapeutic development has been limited by disease rarity and the absence of *in vivo* systems that faithfully capture patient mutations and circuit-level pathology. In this study, we address these limitations by generating a knock-in mouse model harboring the recurrent *TECPR2* c.1319delC mutation and using it to define disease mechanisms, identify translatable functional readouts, and evaluate early gene replacement therapy.

By introducing the patient-derived frameshift mutation into the endogenous mouse locus, we generated a model that reproduces the predominant loss-of-function mechanism observed in affected individuals. Unlike prior knockout or spontaneous mutation models, this knock-in approach preserves the native genomic context while recapitulating the precise pathogenic allele found in patients, increasing its relevance for both mechanistic and translational studies.

Longitudinal phenotyping revealed that homozygous *TECPR2*-KI mice develop reduced body weight, increased grooming behavior, progressive gait abnormalities, and selective tactile sensory impairment, some of which overlap with clinical features described in *TECPR2*-related disorder (5,15). These phenotypes evolve over time and can be quantified reproducibly, establishing the *TECPR2*-KI mouse as a robust platform for tracking disease progression and therapeutic response. A key insight enabled by this model is the identification of selective brainstem circuit vulnerability as a prominent feature of TECPR2 deficiency. Despite broad TECPR2 expression, neuropathology in our model was largely restricted to sensory–motor pathways of the brainstem and spinal cord, with progressive accumulation of axonal spheroids in the gracile and cuneate nuclei and relative sparing of cortical neurons. These findings are consistent with recent work using a neuropathy-associated *Tecpr2* knock-in mouse, which revealed endolysosomal loss-of-function phenotypes in neurons and microglia, supporting a central role for disrupted vesicular and degradative pathways in *TECPR2*-related disease (16). Together, these studies support a model in which TECPR2 loss leads to selective vulnerability of long-range projection neurons and associated circuits, rather than global neurodegeneration.

A key insight enabled by this model is the identification of selective brainstem circuit vulnerability as a central feature of TECPR2 deficiency. Despite broad TECPR2 expression, neuropathology was strikingly restricted to sensory–motor pathways of the brainstem and spinal cord, with progressive accumulation of axonal spheroids in the gracile and cuneate nuclei and relative sparing of cortical neurons. This pattern mirrors neuropathological findings in patients and other TECPR2-deficient species and supports a circuit-selective disease mechanism rather than widespread neurodegeneration. Long-range projection neurons in these pathways likely exhibit heightened vulnerability due to their exceptional reliance on efficient autophagy to maintain axonal homeostasis.

Consistent with this interpretation, ultrastructural and molecular analyses revealed marked defects in autophagic processing within affected brainstem nuclei. Accumulation of abnormal autophagic vacuoles and increased SQSTM1/p62 indicate impaired autophagosome maturation and clearance, directly linking TECPR2 loss to defective autophagy and axonal degeneration. These findings clarify how disruption of a core cellular homeostatic pathway translates into focal circuit dysfunction and highlight autophagy as a mechanistically grounded therapeutic target.

Beyond previously recognized phenotypes, *TECPR2*-KI mice exhibited a pronounced reduction in acoustic startle responses, a behavioral deficit not previously reported in *TECPR2*-related disorder. The acoustic startle reflex depends on well-defined auditory–motor brainstem circuits, and its impairment provides functional evidence of brainstem sensorimotor dysfunction. This finding is particularly relevant given the prominence of brainstem-related clinical features in patients, including impaired reflexes, dysphagia, and central apnea. Importantly, the acoustic startle response represents a non-invasive, quantifiable, and cross-species functional readout, making it a promising biomarker for disease progression and therapeutic efficacy in future clinical studies.

Using this knock-in mouse model, we next evaluated the effects of early gene replacement. Neonatal intracisternal delivery of AAV9-*TECPR2* resulted in widespread transduction of brainstem nuclei and was associated with improvements in autophagy-related markers, a partial reduction in axonal spheroid burden, and improvements across multiple functional outcomes.

Mechanosensory thresholds, gait abnormalities, and acoustic startle responses similarly showed improvement, indicating that early restoration of TECPR2 expression can ameliorate aspects of brainstem-associated dysfunction. These results provide *in vivo* evidence that aspects of TECPR2-related pathology can be modified by postnatal gene replacement. Not all phenotypes were fully prevented. Reduced body weight and excessive grooming persisted in treated mice at later ages, suggesting contributions from early developmental alterations, incomplete vector distribution, or non–cell-autonomous effects involving peripheral or systemic tissues. These findings underscore the importance of therapeutic timing and raise the possibility that even earlier intervention may be required for full disease modification. They also suggest that combination strategies, such as gene replacement paired with pharmacological modulation of lysosomal or autophagy-related pathways, may further enhance therapeutic efficacy.

More broadly, this work informs understanding of selective vulnerability in neurodevelopmental and neurodegenerative disorders linked to autophagy dysfunction. The relative sparing of cortical neurons despite widespread TECPR2 expression highlights how circuit architecture and metabolic demand shape disease expression. By linking impaired autophagic flux to degeneration of specific brainstem sensory–motor pathways, our findings suggest convergent mechanisms that may extend to other disorders characterized by axonal spheroids and brainstem involvement.

Few limitations should be considered. Our analyses focused primarily on the brainstem and spinal cord, and future studies should evaluate peripheral nerves, forebrain circuits, and systemic metabolic contributions. Only a single AAV dose and delivery route were tested, and comparative studies using alternative delivery strategies will be important for optimizing translational approaches. Finally, while the acoustic startle deficit emerges as a promising biomarker, its relevance in patients remains to be established through targeted clinical evaluation.

In summary, we developed a knock-in mouse model carrying the recurrent c.1319delC founder mutation that reproduces the underlying genetic alteration and exhibits circuit-selective pathology and functional impairments associated with TECPR2 deficiency. This model enabled the identification of brainstem sensory–motor circuits as a prominent site of vulnerability and revealed an acoustic startle phenotype that may serve as a translatable functional readout. In addition, early AAV9-mediated TECPR2 delivery was associated with improvement of autophagy-related markers and partial amelioration of select behavioral outcomes. By combining a genetically defined mouse model with quantifiable functional endpoints and a clinically established gene delivery strategy, this study provides a preclinical framework to evaluate TECPR2 gene replacement and to inform future investigations of therapeutic timing, target engagement, and outcome measures.

## MATERIALS AND METHODS

### Generation of the TECPR2 Knock-In Mouse Model

*TECPR2* knock-in (KI) mice were generated at the Harvard Genome Modification Facility. CRISPR/Cas9/guide RNA and donor oligo were injected into fertilized C57BL/6 eggs to generate the equivalent HSP50 exon 8 mutation in the mice. This mutation is a deletion of a single C nucleotide in mouse exon 8: NM_001081057.2; c.1319delC, p.Ser440Serfs19*. Male F1 heterozygous mice were transferred to Charles River Laboratories, where they used these mice for IVF of oocytes from female C57BL/6 to generate heterozygous breeders and used to set up a breeding colony. Mice were genotyped using PCR and sequencing by the Molecular Genetics Core Facility at Boston Children’s Hospital to confirm the presence of the mutation. TECPR2 KI mice were housed in standard clear plastic cages with no more than 5 animals per cage under controlled conditions (lights on 07:00-19:00; humidity 30%-50%; temperature 22-23°C) with ad libitum access to food and water. All experiments were performed between 9:00 and 17:00 in a room maintained at a temperature of 21 ± 1°C. All experimental protocols comply with relevant ethical regulations and were approved by Boston Children’s Hospital Institutional Animal Care and Use Committee (protocol 00002125). Animals were randomized to treatment groups.

### AAV production

Through the Luke Heller TECPR2 Foundation, we have obtained WT human TECPR2and GFP-AAV9 constructs, generated at the Horae Gene Therapy Center, UMass Chan Medical School (Worcester, MA, USA) under the direction of Dr. Guangping Gao. The AAV genome is composed of U1a promoter, TECPR2 CDS, a synthetic short polyA and AAV2 ITRs at both ends. The genome was packaged into the AAV9 capsid by triple-transfection in HEK293 cells.

### In vivo injection of viral vectors

For AAV9 delivery into the CSF via cisterna magna, mice at postnatal day 1–3 (P1–P3) were anesthetized by rapid induction of hypothermia for 2–4 min on ice water until loss of consciousness. The injection site was disinfected with 70% ethanol. A 33-gauge needle attached to a Hamilton syringe was lowered approximately 2 mm into the cisterna magna at a 45° angle, and 1 µL of viral solution (VCAV-04485, 5.00E+12 GC/mL) was slowly injected. Body temperature was gradually restored and maintained on a 37 °C warming pad for 10 min after injection before returning pups to the parental cage. Injections were performed prior to genotyping and blinded to genotype; all pups within each litter received the same viral vector.

### Body Weight and Growth Curve Analysis

Body weights of the TECPR2 KI mice and their WT littermates were recorded with an electronic scale at postnatal day 30, 60 and 90 (P30, P60 and P90, respectively). Growth curves were plotted to compare the development of mutant and WT mice.

### Mechanical Sensitivity Testing

Mechanical sensitivity was assessed using the von Frey filament test. Mice were placed in individual chambers on a mesh platform, and calibrated filaments were applied to the plantar surface of the left hind paw, with a positive response consisting of a paw lifting or flinching response to the fiber. The response patterns were collected and converted into corresponding 50% withdrawal thresholds using the Up–Down Reader software and associated protocol (17). All tests were performed by investigators blinded to the genotype and treatment.

### Acoustic startle reflexes

Mice were tested for startle reflexes in response to broadband auditory stimulation at varying intensities as previously described (18). The animals were tested in a sound shielded startle booth (Kinder Scientific) and the force generated by foot movement was sensed by piezoelectric motion sensor fixed beneath an elevated platform. Mice were placed in a smaller sub chamber anchored to the topside of the platform, which restricted them from rearing on their limbs but freely permitted penetration of the sound stimulus. Sound stimuli were calibrated with the door closed using a sound pressure level meter (Allied Electronics) with the microphone mounted in the position normally occupied by the animal holder. The generation of sound stimuli and recording of force amplitude signals was performed by startle monitor software (Kinder Scientific). Broadband white noise was presented at a background level of 60 dB throughout the experiment and auditory test stimuli consisted of 50-ms broadband white noise pulses in 10-dB steps from 60 to 120 dB. Different intensities of the test stimulus were presented in a pseudo-random order at randomized inter trial intervals (ITI) that varied between 8 and 22 s, with no ITI repeated more than three times. Five repetitions were averaged for each of the intensities for one test subject and responses were normalized to the weight of the mouse tested. Startle response measurements were performed by investigators blinded to the genotype.

### Intoeing and grooming behaviors

Intoeing was quantified using a novel bottom-up imaging and data acquisition technology and analysis platform that provides automated, quantitative, and objective measures of naturalistic rodent behavior in an observer-independent and unbiased fashion as previously described (19). The location of the paws was extracted from body frames using the DeepLabCut and the distance between hindpaws (measure of intoeing) in the unit of pixel distance was calculated. Grooming behaviors were calculated by the time the animal spent grooming the face and body in seconds for 10 minutes recording using the ARBEL algorithm, a machine learning tool with light-based image analysis for automatic classification of 3D behaviors (20).

### Tissue processing and histopathological analysis

Animals were anesthetized with pentobarbital and perfused with a 4% paraformaldehyde (PFA) solution in PBS. The brain was dissected and fixed in 4% PFA solution overnight. Samples were then transferred to a 30% sucrose solution until the tissue sank. Samples were immediately frozen in OCT embedding medium and kept in -80oC until further processing. Coronal cryosections (50 µm) were collected through the brainstem and spinal cord for immunostaining and histopathological analysis. Slices were then stained with hematoxylin and eosin (H&E) to identify spheroids or probed with antibodies for neurofilament and autophagic flux. Quantification of spheroids was done measuring the density of spheroids per mm2 in the cuneate and gracile nuclei areas bilaterally in 10-12 sections per animal, then an average density was calculated using QuPath software (21).

### Immunofluorescence staining

Sections were washed in PBS, permeabilized with PBS + 0.3% Triton X-100 (PBST-0.3) for 15 min and blocked in PBST-0.3 containing 5% normal goat serum (NGS) for 1 h at room temperature. Samples were then incubated overnight at 4 °C with primary antibodies in blocking buffer (Mouse anti-SQSTM1/p62; 1:500; Cell Signaling Technologies #88588; rabbit anti-neurofilament heavy chain; 1:1,000; Sigma Aldrich #N4142). After washes in PBST-0.3, sections were incubated with secondary antibodies for 2 h at room temperature (1:500; Alexa Fluor– conjugated goat anti-rabbit or goat anti-mouse, Invitrogen). Nuclei were counterstained with DAPI (0.5 µg/mL). Sections were mounted with ProLong Glass antifade reagent and imaged on a Zeiss LSM880 confocal microscope. Neurofilament (NF-H) intensity and SQSTM1/p62+ puncta were measured from epifluorescent images. NF-H intensity was measured using sections from WT tissues to set a threshold and the mean pixel intensity from WT and TECPR2 KI tissues was measured using Zeiss Zen 3.8 Software. The quantification of SQSTM1/p62 puncta was done by measuring the area of the cuneate and gracile from epifluorescent images and manually detecting SQSTM1/p62+ puncta in those regions using Zeiss Zen 3.8 Software. Quantitative analysis was performed by an investigator who was blind to the experimental conditions.

### Electron microscopy

Animals were anesthetized with pentobarbital and perfused with 2.5% PFA, 2.5% glutaraldehyde, in 0.1 M cacodylate buffer pH 7.4. Brain samples were fixed with 2.5% PFA, 2.5% glutaraldehyde, in 0.1 M cacodylate buffer containing 5 mM CaCl2 (pH 7.4), post-fixed in 1% osmium tetroxide supplemented with 0.5% potassium hexacyanoferrate trihydrate and potassium dichromate in 0.1 M cacodylate for 1 h, stained with 2% uranyl acetate in double-distilled water for 1 h, dehydrated in graded ethanol solutions, and embedded in epoxy resin (Agar Scientific, AGR1030). Ultrathin sections (70–90 nm) of brainstem, obtained with a LeicaUltracut UCT microtome, were stained with lead citrate and then examined using either a Phillips CM-12 transmission electron microscope (TEM) equipped with a Gatan One View camera or an FEI Tecnai SPIRIT TEM equipped with a bottom-mounted 2 k × 2 k FEI Eagle CCD camera.

### Statistical Analysis

Data are expressed as the mean ± SEM. Significant differences between groups were assessed with one or two-way ANOVA; once the significance of the group differences (p ≤ 0.05) was established, appropriate post-hoc comparisons between pairs of groups were used. Statistical analyses were done using GraphPad Prism Software.

## List of Supplementary Materials

Materials and Methods

Fig. S1 to S3.

## Acknowledgments

Luke Heller TECPR2 Foundation

Jordan Avi Ogman Foundation

Quiver Bioscience [formerly Q-State Biosciences]

Animal Behavior and Physiology Core at Boston Children’s Hospital

Molecular Genetics Core Facility at Boston Children’s Hospital

Cellular Imaging Core at Boston Children’s Hospital

Neurobiology Imaging Facility at Harvard Medical School

Electron Microscopy Core at Harvard Medical School

## Author contributions

Conceptualization: BLT, NH, NA, MF

Methodology: BLT, SL, ASK, NH, NA, MF

Investigation: BLT, PCC, LM, TB, JZ, FV, EC, MA, BZ, SL, NH, NA

Funding acquisition: NH, NA, MF

Project administration: BLT, NH, NA, MF

Supervision: BLT, NA, NH, MF

Writing – original draft: BLT

Writing – review & editing: BLT, NH, ASK, NA, DEF, MF

## Supplementary

**Supplementary Fig. 1:**
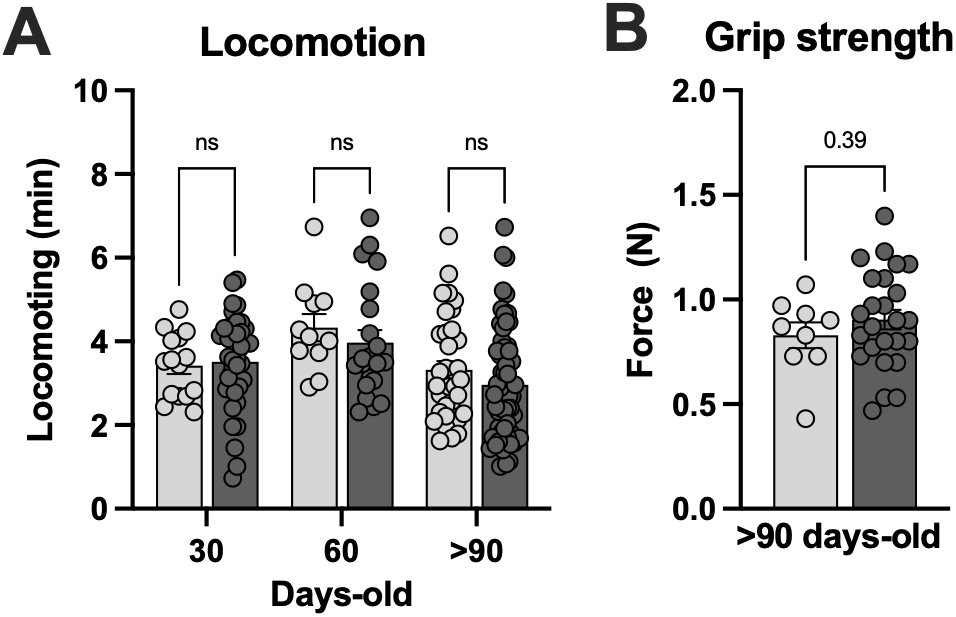
TECPR2-KI mutation [NM_001081057.2; c.1319delC, p.Ser440Serfs19*] does not compromise locomotion and grip strength. **A**, Locomoting time of TECPR2-KI and WT littermate male and female mice at P30, P60 and P90 (n = 5-59 mice per group, data are presented as mean values ± s.e.m., P values from two-way ANOVA followed by Šídák’s multiple comparisons test). **B**, Grip strength of TECPR2-KI and WT littermate male and female mice at >P90 (n = 23-41 mice per group; data are presented as mean values ± s.e.m., P values from unpaired t test). C, Mechanical withdrawal thresholds of TECPR2-KI and WT littermate male and female mice using an up–down von Frey method at >P90 (n = 9-25 mice per group; data are presented as mean values ± s.e.m., P values from unpaired t test).

**Supplementary Fig. 2:**
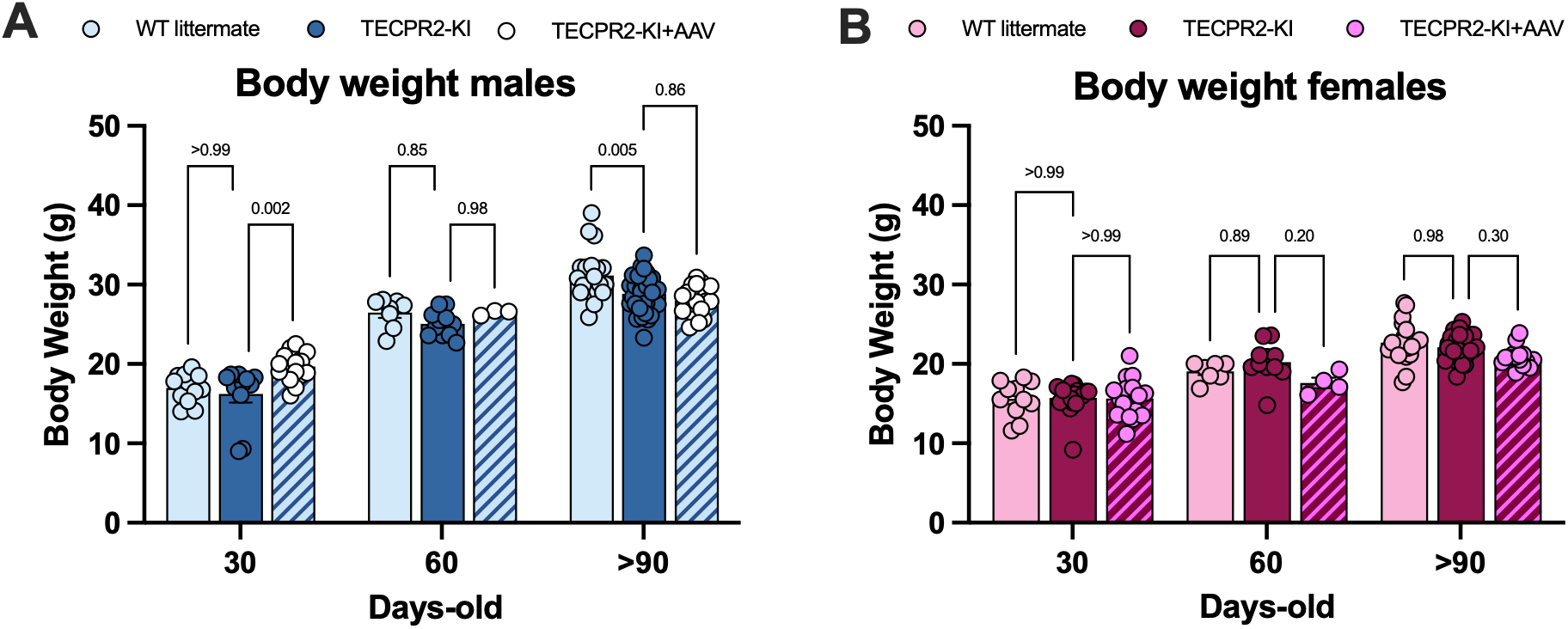
TECPR2-KI mutation [NM_001081057.2; c.1319delC, p.Ser440Serfs19*] induces changes in body weight in male mice. **A**, Body weight of TECPR2-KI and WT littermate male and **B**, female mice at P30, P60 and P90 (n = 4-33 mice per group, data are presented as mean values ± s.e.m., P values from two-way ANOVA followed by Šídák’s multiple comparisons test).

**Supplementary Fig. 3:**
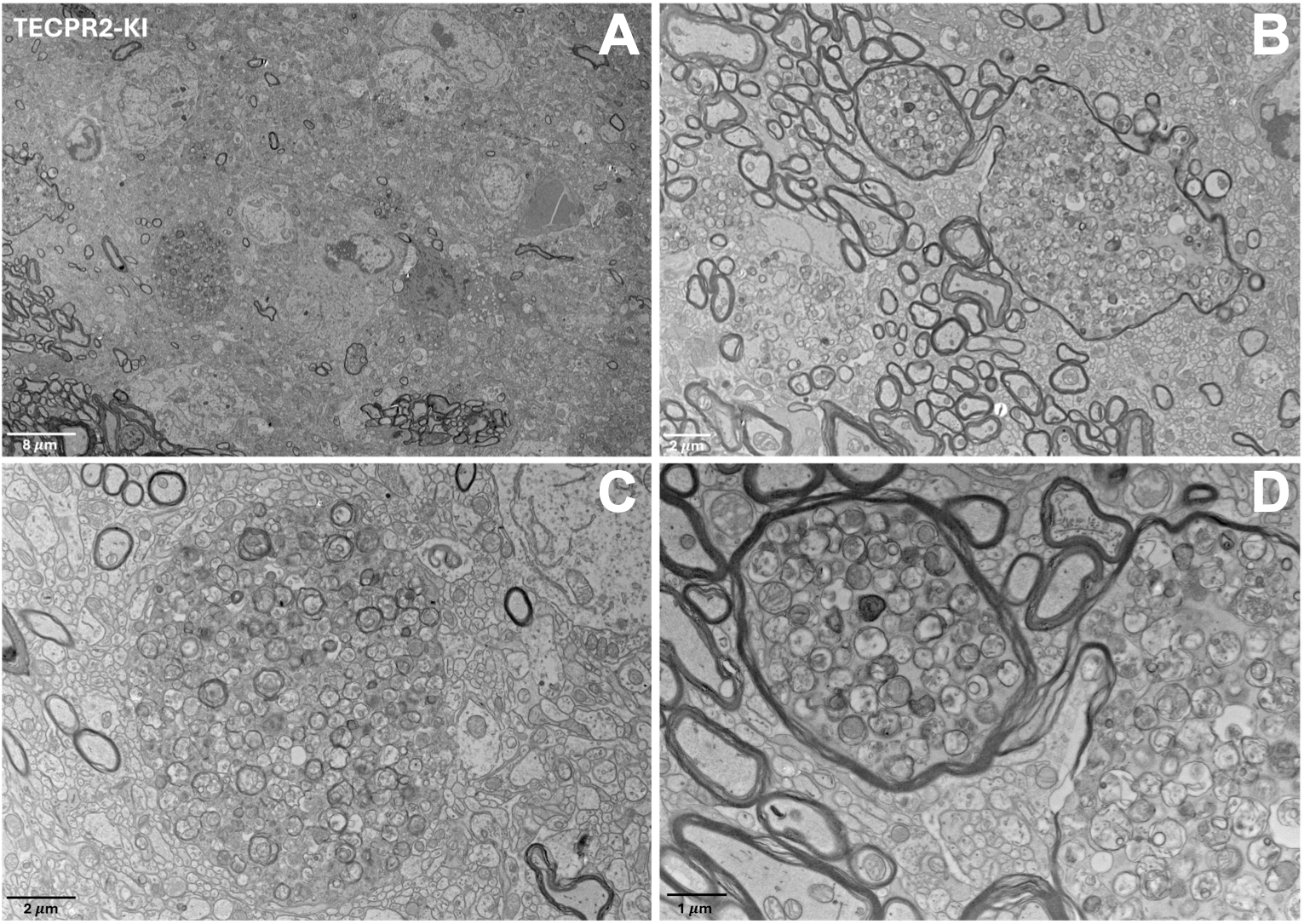
TECPR2-KI mutation induces structural abnormalities in TECPR2-KI mouse brainstem. **A-D**, Ultrastructural analysis by transmission electron microscopy revealed prominent structural abnormalities in TECPR2-KI mouse brainstem.

