## supplementary for "Preclinical validation of AAV9-TECPR2 gene therapy in a novel knock-in model of TECPR2-related disorder"

**Abstract**

*TECPR2*-related disorder is a rare, autosomal recessive neurodevelopmental and neurodegenerative disease characterized by early-onset motor dysfunction, sensory- and autonomic neuropathy, and progressive neurological decline with early mortality. Currently, there are no effective treatments for individuals affected by this debilitating condition. To advance our understanding of disease mechanisms and explore therapeutic strategies, we developed and then characterized a knock-in (KI) mouse model carrying the human *TECPR2* c.1319delC frameshift mutation.

TECPR2-KI mice exhibit a subset of disease-relevant phenotypes, most prominently abnormal gait, along with reduced body weight and altered tactile sensitivity. We additionally observe a reduction in acoustic startle responses, consistent with dysfunction of brainstem-associated sensorimotor pathways. Histopathological analyses reveal progressive accumulation of axonal spheroids in the dorsal column nuclei, together with abnormalities in autophagy-related markers, features previously reported in individuals with *TECPR2*-related disorder.

To assess the therapeutic potential of gene replacement, we delivered *TECPR2* via intracisternal infusion of AAV9/*TECPR2* in neonatal KI mice. Gene therapy restored mechanosensory function, normalized gait and startle responses, maintain autophagic homeostasis, and partially reduced axonal pathology. These findings demonstrate that *TECPR2*-associated deficits are not only replicable in this new mouse model but are also amenable to postnatal intervention.

Our study introduces a genetically accurate murine model of TECPR2 deficiency, identifies brainstem-associated phenotypes, and provides preliminary evidence supporting the feasibility of AAV9-mediated TECPR2 gene delivery, establishing a foundation for future translational research in a currently untreatable disease.

### Supplementary

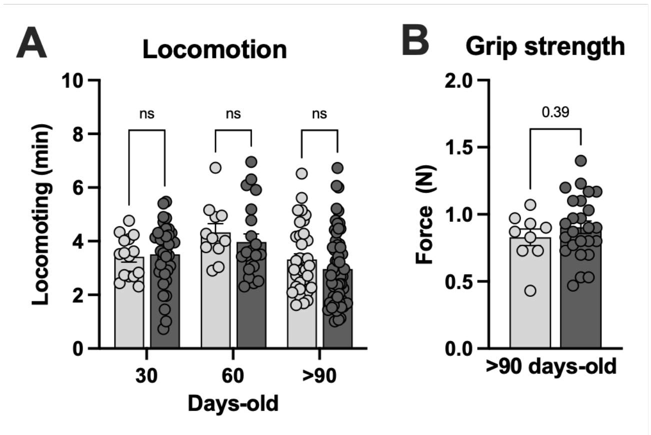


**Supplementary Fig. 1**: ***TECPR2*-KI mutation [NM_001081057.2; c.1319delC, p.Ser440Serfs19*] does not compromise locomotion and grip strength. A,** Locomoting time of *TECPR2*-KI and WT littermate male and female mice at P30, P60 and P90 (*n* = 5-59 mice per group, data are presented as mean values ± s.e.m., *P* values from two-way ANOVA followed by Šídák's multiple comparisons test). **B,** Grip strength of *TECPR2*-KI and WT littermate male and female mice at >P90 (*n* = 23-41 mice per group; data are presented as mean values ± s.e.m., *P* values from unpaired t test). **C,** Mechanical withdrawal thresholds of *TECPR2*-KI and WT littermate male and female mice using an up–down von Frey method at >P90 (*n* = 9-25 mice per group; data are presented as mean values ± s.e.m., *P* values from unpaired t test).

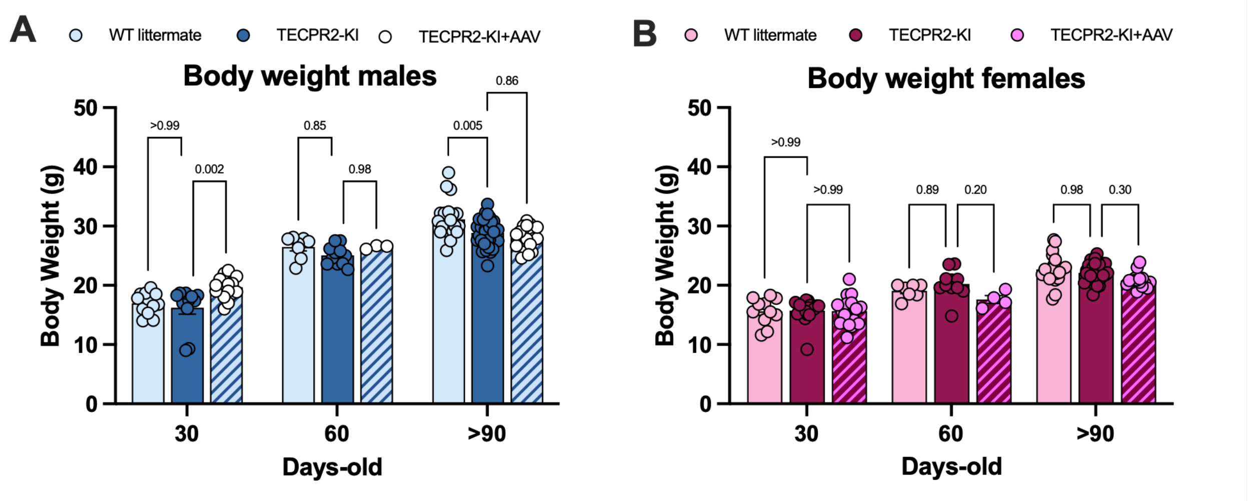


**Supplementary Fig. 2**: ***TECPR2*-KI mutation [NM_001081057.2; c.1319delC, p.Ser440Serfs19*] induces changes in body weight in male mice. A,** Body weight of *TECPR2*-KI and WT littermate male and **B,** female mice at P30, P60 and P90 (*n* = 4-33 mice per group, data are presented as mean values ± s.e.m., *P* values from two-way ANOVA followed by Šídák's multiple comparisons test).

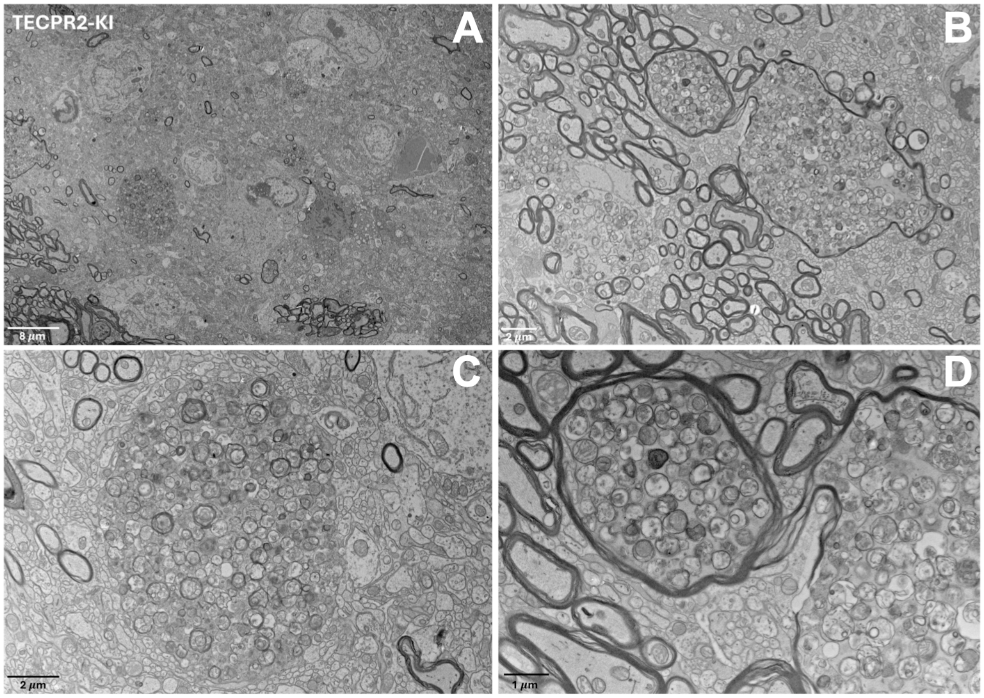


**Supplementary Fig. 3**: ***TECPR2*-KI mutation induces structural abnormalities in *TECPR2*-KI mouse brainstem. A-D.** Ultrastructural analysis by transmission electron microscopy revealed prominent structural abnormalities in *TECPR2*-KI mouse brainstem.
